# Screening for the ancient polar bear mitochondrial genome reveals low integration of mitochondrial pseudogenes (*numts*) in bears

**DOI:** 10.1101/094771

**Authors:** Fritjof Lammers, Axel Janke, Cornelia Rüecklé, Vera Zizka, Maria Nilsson

## Abstract

Phylogenetic analyses of nuclear and mitochondrial genomes have shown that polar bears captured the mitochondrial genome of brown bears some 160,00 years ago. This hybridization event likely led to an extinction of the original polar bear mitochondrial genome. However, parts of the mitochondrial DNA occasionally integrates into the nuclear genome, forming pseudogenes called *numts* (nuclear mitochondrial integrations). Screening the polar bear genome for *numts,* we identified only 13 such integrations. Analyses of whole-genome sequences from additional polar bears, brown and American black bears as well as the giant panda indicates that the discovered *numts* entered the bear lineage before the initial ursid radiation some 14 million years ago. Our findings suggests a low integration rate of *numts* in the bear lineage and a complete loss of the original polar bear mitochondrial genome.

---

Polar and brown bears are two well-recognized species, that differ in their morphology and ecology (Nowak 1999). Recent research has shown that polar bears diverged from brown bears in the mid Pleistocene (Hailer et al. 2012; Liu et al. 2014), which is supported by the fossil record (Kurtén and Anderson 1980) (but see Miller et al. (2012), for earlier divergence estimates). Phylogenetic analyses of mitochondrial DNA (mtDNA) showed that polar bears appear to be nested inside the brown bear radiation and dated the emergence of polar bears around 160 thousand years ago (kya) (Hailer et al. 2012; Edwards et al. 2011; Lindqvist et al. 2010). The deviating mtDNA phylogeny can be explained by recurrent introgressive hybridization between female brown bears and male polar bears 160 kya resulting in a mitochondrial capture event that replaced the original polar bear mitochondrial (mt) genome (Edwards et al. 2011; Hailer et al. 2012; Miller et al. 2012a). As evident from the low genetic diversity among polar bears, population bottlenecks during interglacials led to a fixation of the introgressed brown bear mtDNA in the polar bear lineage. Consequently, the original polar bear mtDNA was lost in all extant populations of polar bears.

Occasionally mt genome sequences are transferred to the nucleus, and become incorporated as pseudogenes called *numts* (**nu**clear sequence of **mit**ochondrial origin, pronounced “new-mite”) (Hazkani-Covo, Zeller, and Martin 2010; Tsuji et al. 2012; Lopez et al. 1994; Rogers and Griffiths-Jones 2012; Dayama et al. 2014). *Numt* insertions occur via non-homologous end joining at double strand breaks in the nuclear genome. In general, the genomic fraction of *numts* is less than 0.1 % with the highest proportion found in plants and yeast (Kolokotronis, Macphee, and Greenwood 2007; Dayama et al. 2014). Several genome scale studies have discovered that the total copy number and sequence length of *numts* varies widely between mammalian species (Hazkani-Covo, Zeller, and Martin 2010). For instance only 49 copies, totalling 6 kilo base pairs (kb), are found in the rat (*Rattus norvegicus*) genome while 1859 copies (2,093 kb) are found in opossum (*Monodelphis domestica*) (Hazkani-Covo, Zeller, and Martin 2010). Three different processes can contribute to the differences in the number of *numts* between species: the frequency of mitochondrial transfer, the amount of integrations as well as the dynamics of insertion processes.

To date *numt* insertions have not been studied in representatives of the bear family (Ursidae). If indeed, at 160 kya, the polar bear mt genome was replaced by the brown bear mt genome, all polar bear *numts* that entered the nuclear genome between the divergence of both species (~600 kya) and the mitochondrial capture event (~160 kya) are genomic ‘fossils’ of the original polar bear mtDNA (Figure 1). We screened the polar bear genome for *numts* to reconstruct the ancient polar bear mt genome sequence. [Position Figure 1] Although the rate of *numt* integration in the genome is generally low, polymorphic *numt* copies are known from the human population (Dayama et al. 2014) indicating ongoing *numt* integration even within a few hundred generations. Screening the polar bear genome sequence, using the mt genome found in extant polar bears, identified 64 putative *numts* totalling about 29 kb sequence (Table 1). The identified *numt* sequences cover 62% of a bear mt genome, i.e. 38% of the mt genome did not contribute to the polar bear *numt* landscape. We focused our analyses on 22 *numts* that were longer than 200 bp, which represent 57% of the mt genome and originate mainly from the NADH1 to COII region (Figure 2). Both rRNA genes as well as NADH4 and NADH5 are in addition partially covered. [Position Figure 2] Phylogenetic maximum likelihood analysis of the *numt* sequences, including homologous mitochondrial sequences of all living bear species and other carnivores, indicated an old age of the *numts* (Supplementary Material Data 1: Phylogenetic trees). Fourteen of the 22 identified *numts* are situated in close proximity to each other in the nuclear genome and the gene order and orientation suggest that these are eroded fragments from single ancient integrations. Merging the consecutive fragments reduced the total number to 13 *numts* (Supplementary Table S2). Aligning the genomic polar bear *numt* loci to orthologous sequences from the giant panda (*Ailuropoda melanoleuca*) revealed that 11 of the 13 *numts* are present to full extent in the giant panda genome and thus integrated at least 14 million years ago when the giant panda lineage diverged (Kumar et al. 2016). Locus 7 was only partially identified in the giant panda genome and locus 8 was not recovered. The genomic distance between *numt* fragments in the polar bear genome, matched approximately the distance of the same fragments when mapped to the mt genome (Supplementary Table 3). The distance difference between *numt* fragments in the nuclear genome their mitochondrial origins can be explained by integration of transposable elements (loci 7, 9 and 10) or expansion of short tandem repeats (loci 6 and 11). In locus 10, a full LINE1-1_Ame transposable element inserted in the giant panda genome. Nearly all identified polar bear *numt* sequences seem to have inserted prior to the evolution of Ursidae. Consequently, no *numts* representing ancestral or recent polar bear mtDNA have entered the nuclear genome (Figure 1). As an additional line of evidence, we analyzed whole-genome sequencing data of two additional bear species for polymorphic *numt* insertions among Ursidae using the structural variation (SV) caller Lumpy (Layer et al. 2014). If *numts* present in the polar bear genome integrated after the divergence of a closely related bear species, a genomic deletion matching approximately the size of the *numt* would be present in the other species’ genome. Our SV analyses identified that the *numt* loci in polar bear do not coincide with deletions in the genomes for brown bear ( *Ursus arctos)* or American black bear (*Ursus americanus*) (Supplementary Data S2: Excel spreadsheet). These additional findings suggest that the *numts* were at least present since American black bears diverged from the lineage to polar and brown bear, some 3 Mya (Kumar et al. 2016). Thus, the SV analyses support our hypothesis that all numts detected in the polar bear genome sequence occurred before the divergence of extant bears.

**Figure 1.**
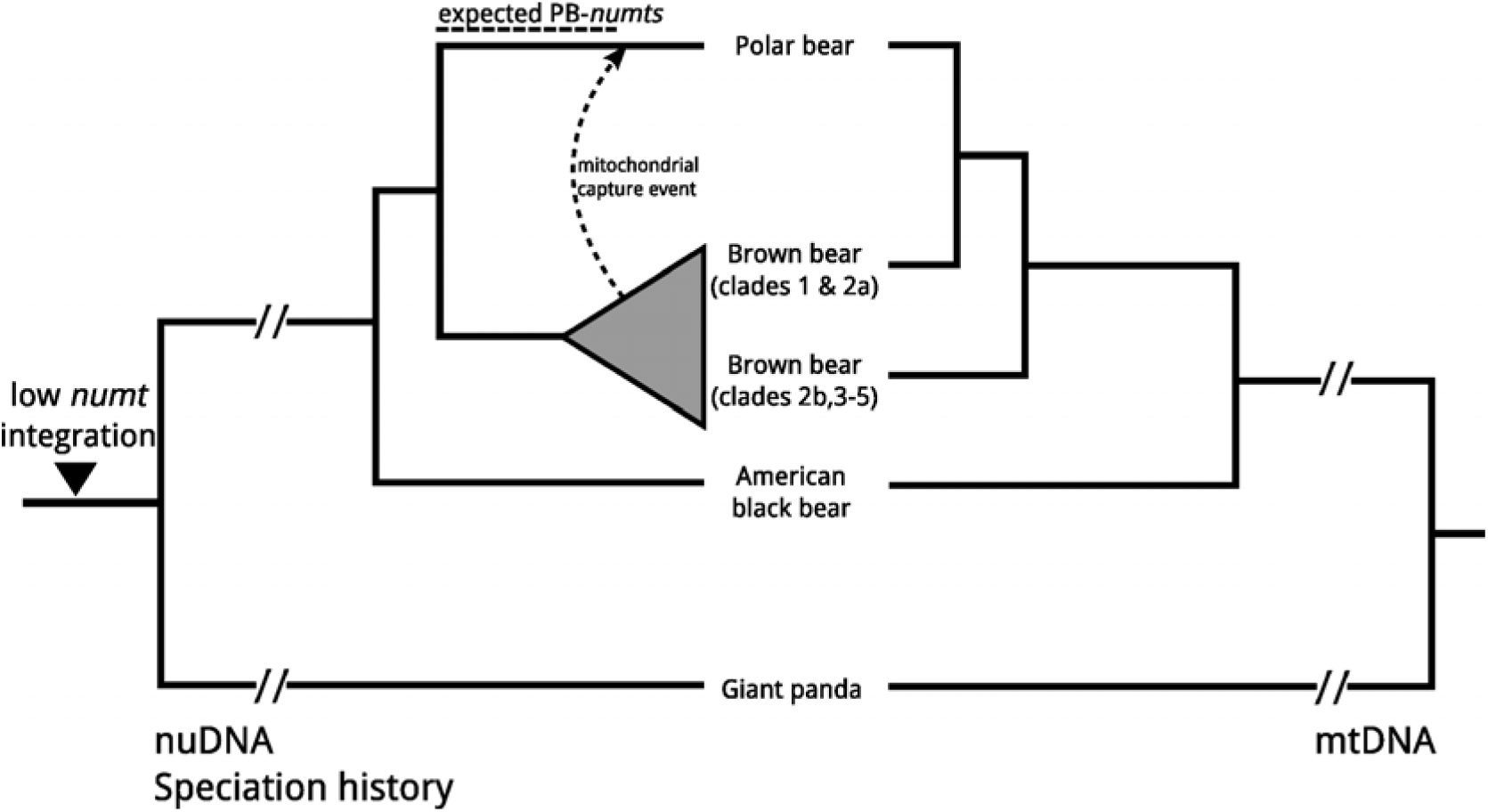
Phylogeny of bears reconstructed by nuDNA (left side) and mtDNA (right side). The left phylogeny reflects the speciation history of bears. About 160 kya, the original polar bear mtDNA lineage was replaced by brown bears (dashed arrow) causing the observed paraphyly of brown bears in the mtDNA phylogeny (right side). Dashed lines above the nuDNA phylogeny indicate the timeframe for potential integration of *numts* that represent the original polar bear mtDNA (PB-numts) and the observed reduction of *numt* integration in Ursidae.

**Table 1.**
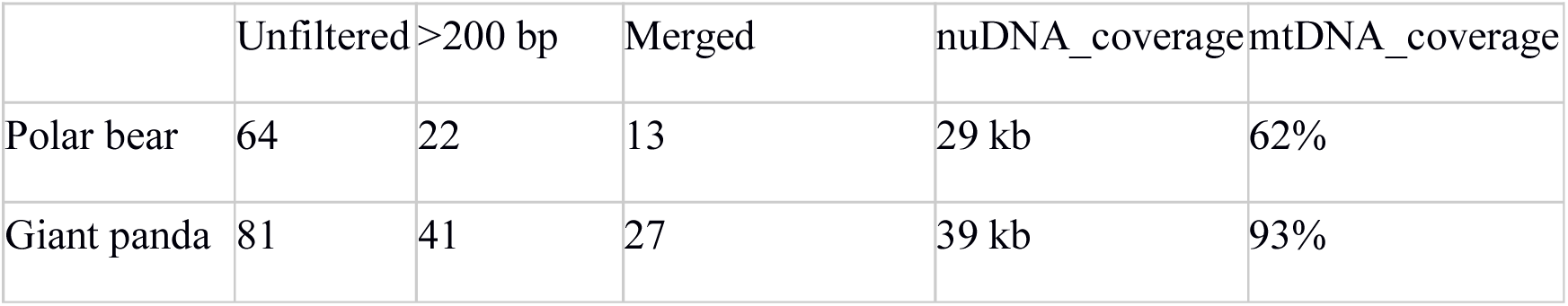
*Numts* identified in polar bear and giant panda. Shown are the unfiltered number of BLAST hits in the respective genome, the number of *numts* that is longer than 200 bp, and the number of *numts* after merging those within 10 kb distance to each other. Additionally shown are the amount of nuclear genome sequence covered and the percentage of mtDNA covered by unfiltered *numts*.

**Figure 2.**
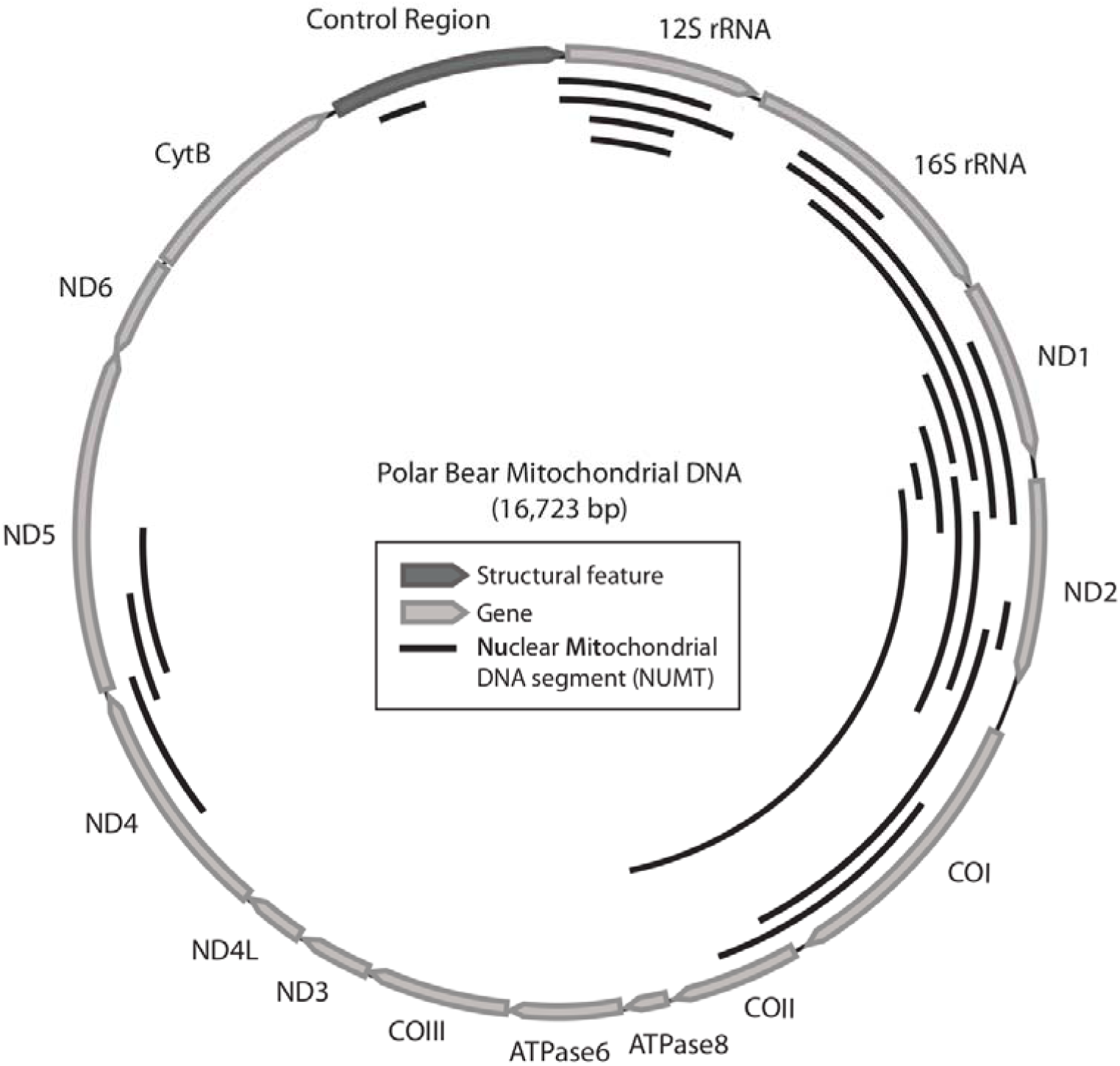
Genetic map of the polar bear mitochondrial genome with annotated genes. The identified *numts* longer than 200 bp are shown as solid black lines on the inside of the mt genome. For exact position and labelling of *numt* sequence refer to Supplementary Table 2.

It is not unreasonable, that the number of *numts* and the genomic fraction derived from *numts* appear low in the polar bear genome when compared to other mammals (Hazkani-Covo, Zeller, and Martin 2010). The two murid rodents, mouse (*Mus musculus*) and rat only have between 6-39 kb numts in their respective genomes, which is among the lowest numbers of *numts* in mammals (Hazkani-Covo, Zeller, and Martin 2010). The low incidence of *numts* in the closely related mouse and rat, as well as in the bear family suggest that these groups have evolved mechanisms to suppress the integration of numts or that the rate of deletion is higher than in other groups (Hazkani-Covo, Zeller, and Martin 2010).

However, another important reason for the lack of recent *numts* in the polar bear genome may lie in the genome assembly processes. Recent *numts* are highly similar to the actual mitochondrial genome, and de Bruijn-graph based assembling algorithms might not be able to distinguish between short reads originating from *numts* or mtDNA (Li, Zhu, et al. 2010; Hahn, Bachmann, and Chevreux 2013). This would cause a severe under-representation of recent *numts* in short-read based genome assemblies, and may in addition have introduced a bias in many published genome assemblies. So called, third generation sequencing technologies like PacBio or Nanopore produce long reads, that are more likely to span complete *numt* insertion and thus facilitate their incorporation into genome assemblies (Sohn and Nam 2016). To our knowledge, the gorilla genome is the only non-hominid mammalian genome generated by extensive usage of PacBio sequences (Gordon et al. 2016) but we anticipate several additional long-read based genome assemblies to become available in the next years. Long-read based sequencing of a bear genome can give further insights into the fate of *numts* in Ursidae.

Recently inserted *numts* create problems for population and phylogenetic analyses of mt genes, if they are PCR amplified instead of the mt genes (Bensasson et al. 2001), but appear absent from the bear family. Thus, previous mitochondrial studies of bear phylogeny and population structures (Edwards et al. 2011) would be free from artefacts in the form of mitochondrial pseudogenes.

## Conclusions

The current polar bear genome assembly lacks recently inserted *numts.* All identified *numt* insertions are at least 14 My old. A low *numt* insertion frequency has been reported for rodents and might be common in other mammalian groups. Thus, a low number of *numts* in bear genomes is not unreasonable, but an artefact from modern whole genome assembly algorithms cannot be excluded until further *de novo* assembled bear genomes become available. Utilizing long-read high-throughput technologies like PacBio or Nanopore might yield further insight into the fate of *numts* in Ursidae or other taxonomic groups with suspected *numt* depletion.

## Materials and Methods

### BLAST search for numts in the polar bear genome

The mitochondrial genome sequence of the polar bear (AJ428577.1) was screened against the polar bear genome sequence (Liu et al. 2014) using BLAST (Altschul et al. 1990) with a word size of 20 bp to identify and localize insertions of mitochondrial DNA (*numts*) in the genome. The identified *numts* were filtered for length, and 22 *numt* hits longer than 200 bp were subjected to further analyses. *Numts* located within 10 kb distance to each other in the nuclear genome were merged to clusters, that consisted of two to three fragments interspersed by genomic DNA.

### Finding numts homologs in mitochondrial genomes of selected carnivores

The *numt* sequences from the polar bear genome were blasted against circularized (i.e. self-concatenated) mt genomes from all bear species as well as dog (*Canis lupus familiaris*) and cat (*Felis catus*) (Supplementary Table 1) using word_size 20 and an Evalue cutoff of 0.01.

### Phylogenetic reconstruction of numt loci

The polar bear *numt* sequences were aligned with mitochondrial homologs of Carnivora using MAFFT v7.305b (Katoh and Standley 2013) applying the --adjustdirection option. Alignments were trimmed with trimal (Capella-Gutiérrez, Silla-Martínez, and Gabaldón 2009) using “automated1” mode. RAxML 8.2.9 (Stamatakis 2014) calculated phylogenetic maximum likelihood trees using the GTRGAMMAI model. Node support was computed with 1000 bootstrap replicates.

### Creating 2-way alignments of polar bear/giant panda and mitochondrial homologs

The clustered *numt* sequences plus 1 kb flanking sequences from the polar bear genome were queried against the giant panda sequence (Li, Fan, et al. 2010) using BLAT (Kent 2002). The list was sorted based on the alignment length and the ratio of BLATSCORE/ALIGNMENTLENGTH and manually screened for the extent of matching sequence between the polar bear *numt* loci and giant panda genomic sequence. The identified orthologs were aligned against polar bear sequences using MAFFT (Katoh and Standley 2013) and manually inspected. The resulting 2-way alignments between polar bear and giant panda, corresponding *numt* sequences from other carnivore species were aligned using AliView (Larsson 2014) and MAFFT. Repetitive element screening were performed on the GIRI webserver (Kohany et al. 2006) and manually included in the alignment.

### Whole-genome structural variation screen for deletion associated with numt insertions in polar bear

For further investigation, whole-genome sequencing reads of two additional polar bear individuals (Accession no. SRR518686, SRR518687 and SRR518661,SRR518662) (Miller et al. 2012a), one brown bear (SRR935592, SRR935595, SRR935624, SRR935628) (Liu et al. 2014) one American black bear (SRR518723) (Miller et al. 2012a) were mapped against the polar bear genome as described elsewhere (Kumar et al. 2016). Lumpy (Layer et al. 2014) screened the genomes for structural variation using default settings. Predicted deletions longer than 200 bp were extracted from the SV dataset and tested for spatial association between genomic deletions of 100 bp and 10 kb in size and *numt* coordinates in the polar bear genome were investigated using bedtools multiinter (Quinlan and Hall 2010). Additionally, the short-read mapping was manually inspected using the Integrative Genome Viewer (Thorvaldsdóttir, Robinson, and Mesirov 2013).

## Acknowledgments

The authors thank S. Gallus for creating the mitochondrial genome annotation figure.

